# Predicting forest damage using relative abundance of multiple deer species and national forest inventory data

**DOI:** 10.1101/2023.08.17.553670

**Authors:** Colin Brock, Virginia Morera-Pujol, Kilian J. Murphy, Maarten Nieuwenhuis, Simone Ciuti

## Abstract

Human modification of landscape and natural resources have facilitated deer population irruptions across the world resulting in widespread human-wildlife conflicts. These conflicts occur across the field of natural resource management and negatively affect both the public and vested stakeholders when their livelihoods are placed at risk, for instance, the forestry sector. Deer, both native and non-native, at high densities can damage forest ecosystems impacting biodiversity and ecological functioning at multiple levels and can inflict large ecological and economic costs. The ecological drivers of forest damage and the roles of single and multiple co-occurring deer species is not well understood due to a lack of coordinated high resolution deer distribution, deer abundance and forest damage data. Here, we aim to disentangle the relationship between forest damage, forest characteristics and the roles deer play in damaging forest ecosystems. To achieve this, we adopt a novel approach integrating recent high resolution deer distribution data for multiple deer species (native and non-native) and combining them with forest inventory data collected in 1,681 sampling stations across Ireland to provide risk scenario predictions for practitioners to use on a national scale. Forest characteristics played a key role in the severity and type of damage risk that deer posed. We found all damage types were more prevalent in forests with greater tree densities where deer are more likely to find refuge from human disturbance. Bark stripping damage was more prevalent in mature forests with high tree diversity and ground level flora (e.g., bryophytes, herbs, and shrubs). Similarly, browsing damage was more prevalent in forests with greater tree richness but with understorey vegetation dominated by grass and ferns. Fraying damage was more common in mixed woodlands with understory dominated by bryophytes and grass. Crucially, we found that type and severity of forest damage were shaped by the interaction of multiple deer species occurring simultaneously, particularly at high densities, suggesting subtle inter-species competition and exclusion/partition dynamics that require further investigation to understand the ecological mechanism. Finally, we produce risk scenarios of forest damage by co-occurring deer species and precisely predict where damage is likely to occur on a national scale. We predict high levels of damage in sika and/or red deer hotspots, matching areas of highly concentrated deer distributions. This study highlights the ecological drivers and the role that co-occurring native and non-native deer species have on forest damage within a large spatial scale. By combining reliable species distribution models with the national forest inventory data, we can now provide a useful tool for practitioners to help alleviate and mitigate forest damage and human wildlife conflicts.

## Introduction

Large herbivores are keystone species within terrestrial ecosystems, and their presence and abundance can influence species composition and ecosystem function (Rooney and Waller, 2003; Bernes et al., 2018). Large herbivore population dynamics are naturally regulated by both abiotic and biotic factors, including food availability, weather conditions, and predator-prey interactions (Bobek, 1977; Melis et al., 2009; Ciuti et al., 2015). Unmodified ecosystems in the Anthropocene are rare, with the encroachment of human infrastructure and activity into wild places. Within natural systems large herbivores play a beneficial role in ecosystem productivity (Frank and McNaughton, 1993; Augustine et al., 2003; van Klink et al., 2014, Bakker et al., 2015), shape plant communities by promoting community assembly and species coexistence in reducing competition among floral species (Nishizawa et al., 2016), increase plant biodiversity via seed dispersal, and directly create invertebrate habitats via faecal droppings (Purser, 2009). In natural ecosystems with predator-prey interactions and trophic cascades in place, predators shape large herbivores survival, reproduction, and overall population dynamics through direct and indirect mortality by creating a landscape of fear that increases stress and vigilance in prey species via top-down effects (Brown and Kotler, 2004; Schmitz et al., 2004; Preisser et al., 2005; Zanette and Clinchy, 2020; Palmer et al., 2022). Thus, within human-unmodified landscapes, native large herbivores have evolved in the natural ecosystem they live in, occupy a specific trophic niche and are an important component of our biodiversity (Augustine et al., 2003; van Klink et al., 2014; Bakker et al., 2015).

Where large herbivore populations occur within human-modified landscapes, often where predator density is reduced from an historical baseline, or predators have been eradicated completely, non-natural ecosystems can drive population dynamics through both direct and indirect anthropogenic bottom-up and top-down effects that influence trophic cascades (Dorresteijn et al., 2015), often times resulting in population irruptions that have caused large herbivore species to become prevalent at unsustainable population densities (Côté et al., 2004). This is evident in the literature throughout the globe, where unsustainable population growth in recent decades has led to increased human-wildlife coexistence issues (Côté et al., 2004; Apollonio et al., 2010). Generally, population increases to or beyond carrying capacity can be attributed to agricultural and silvicultural expansion (Alverson, 1988; Porter & Underwood, 1999; Fuller & Gill, 2001), milder winters (Solberg et al., 1999; Loison et al., 1999) and the reduction or removal of large natural predators (Boitani, 1995, Paquet & Corbyn, 2003). The effect of deer in areas with overabundant populations may shift from beneficial to damaging to the overall biodiversity of fauna and flora species. For instance, over-browsing, particularly at high densities, can reduce the abundance and densities of ground nesting birds (Newson et al., 2012), insects (Allombert et al., 2005), soil mesofauna (Katagiri and Hiiji, 2017) and suppress pollinator species (Wilkerson et al., 2013). Damage can extend to agricultural croplands (Conover and Decker, 1991), forest nurseries and orchards (Porter, 1983; Conover, 1984) impacting important socio-economic industries.

In forest environments, damage caused by deer at high densities can inhibit forest regeneration (Tanentzap et al., 2009; Borowski et al., 2021), reduce understory cover and richness, and increase the success rate for invasive plant species (Rodewald & Arcese, 2016). Over browsing can homogenize plant species composition, affect seed and sapling abundance (Catorci et al., 2016; Lessard et al., 2012), decimate understorey foliage (Eichhorn et al., 2017), lower plant fecundity (Knight, 2004), and indirectly affect the reproduction of woody plants (Sakata and Yamasaki, 2015). Deer can impact biological diversity and community structures within forests at every level and have the potential to compromise ecological integrity and stability (e.g., Beguin et al., 2011; Goetsch et al., 2011; Shelton et al., 2014; Nakahama et al., 2020; Borrowski et al., 2021, Bucher et al., 2021). Commercially, ecological instability and reduction in tree growth leads to higher susceptibility to erosion and flooding (Reimoser, 2003), longer stand rotations and lower stand densities (Kullberg & Bergstorm, 2001), ultimately reducing productivity and loss of profit, again, leading to the exacerbation of the conflict with human activities.

Three deer behaviours typically lead to forest damage: bark stripping, browsing, and fraying (Gill, 1992; Barrios-Garcia et al., 2011). Bark stripping can cause severe damage to trees as deer peel off and remove strips of bark from a tree, leaving it exposed and vulnerable to disease (Vasiliauskas, 2001; Kiffner et al., 2008). Browsing is characterized by the tearing of leaves and stalks for consumption. Male adults fray, or rub, their antlers on trees, vegetation, and on the soil to either remove the velvet from their growing antlers or to leave their scent for marking territories and attracting females (Miller et al., 1991; Carranza and Mateos-Quesada, 2001). These behaviours are natural and would be expected in any environment in which deer are present. Issues may arise when the populations of deer increase and ecological heterogeneity is lost, exposing both natural and commercial forests to high levels of damage. The likelihood of damage occurring in a forest is likely to be related to the density of deer inhabiting the area (of either a single species or co-occurring multiple species), although the nature of such a relationship may be quite complex and governed by several other factors (Putman et al., 2011). Intensity of forest damage by deer can be further influenced by factors such as the balance of natural food supply and other “attraction factors” (Reimosur and Gossow, 1996), landscape type or attributes (e.g., afforestation landscapes are more susceptible to damage; Reimoser, 2003; Spake et al., 2020), and regional climate conditions (Spake et al., 2020). It is unlikely there is a single threshold of deer density where damage becomes more likely, but rather a combination of broad density ranges, “attraction factors” and the length of time that over abundant populations have been present in an area (Putman et al., 2011). Putman et al. (2011) argued that assessing the impacts of deer on forests, coupled with accurate estimates of deer densities are the best practice in developing effective management strategies to reduce or mitigate the harmful effects of deer on forests. While much of the literature on forest damage by deer species have focused on a single species in localized areas, there is little known about how multiple species, particularly on a national scale, can influence and drive damage in forests.

Understanding the mechanisms that drive forest damage by deer will ensure effective management practices being implemented to alleviate and mitigate the impacts. To achieve this, it is essential to monitor and collect empirical data on both forests and deer species presence simultaneously. Ireland is an excellent example to examine these impacts as it is home to three established and widespread species of deer, one native and two non-natives (Morera-Pujol et al., 2023). In addition, Ireland has few remaining natural environments and is dominated by extensive agricultural practices, rapid urbanisation, and landscape fragmentation (Boyle, 2009), a landscape that is common worldwide. Ireland differs from other countries however, as forest cover has increased in the past century by nearly 700,000ha, or roughly 10% of the national land area, especially in the past 40 years, with many of these new woodlands lacking diversity as they are primarily single species tree plantations used for commercial forestry (DAFM, 2017). This forest expansion is one of the main drivers behind the population explosion of deer species and the expansion of deer distribution spreading into young, new forests previously uninhabited throughout the country (Carden et al., 2011, Morera-Pujol et al., 2023, Murphy et al. 2023). Recently published, up to date data on the widespread presence of deer (Morera-Pujol et al., 2023) can be combined with the most recent available forest damage data (DAFM, 2017) across the country to provide forest managers and other stakeholders with the knowledge and tools to allow them to prevent damage and alleviate human-wildlife conflicts. The Irish case study can also be used as a blueprint for many other countries experiencing the same level of human-deer conflicts, providing a tool for practitioners to plan into the future.

Here, we take the recommendation by Putman et al. (2011) that the best practice for reducing and mitigating the harmful effects of deer is by developing management strategies founded on the assessment of accurate deer densities and their impacts on forests. We offer a novel insight into the relationship between forest damage and the abundance and distribution of native and non-native deer species, and their relationships with each other and certain forest characteristics. We did this by combining deer damage information from Ireland’s National Forest Inventory (NFI) with the most up to date and comprehensive predicted relative distributions and abundance estimates for three species of deer (red, fallow and sika) on a national scale (Morera-Pujol et al., 2023). For the first time, to our knowledge, we examine these relationships by including three types of forest damage, each occurring through different behavioural processes, with multiple species of deer on a national scale. Our aim is to highlight forest characteristics that are “attractive” for deer to cause damage, understand what deer species are likely to cause damage and introduce a tool to predict risk scenarios that will enable effective management decisions aimed at preventing damage to the national forest estate and alleviate human wildlife conflicts. Specifically, we aim to; i) define the forest composition where specific types of damage are more common; ii) highlight what deer species, or co-occurring species, are causing high levels of forest damage; iii) map predictive damage likelihood for each damage type across the country. We hypothesize that: a) forest composition will differ for each damage type, particularly for browsing damage as the behaviour for bark stripping and fraying is more similar; b) one, or both, of the non-native deer species are expected to causing high levels of damage due to their higher relative abundance (Morera-Pujol et al., 2023); and c) the likelihood of forest damage will be concentrated in deer “hotspot” areas, potentially highlighting the importance for foresters to be constantly updated by wildlife biologists on the status and trends of deer population at different spatial scales.

## Methods

### Data Overview

There are four species of deer in Ireland, three of which are well established i.e., the native red deer (*Cervus elaphus)* and two non-native species, sika deer (*Cervus nippon),* and fallow deer (*Dama dama)* (Carden et al., 2011). The fourth, muntjac deer *(Muntiacus reevesi),* is the most recently introduced species to the island with very limited distribution. The known distribution of muntjac deer does not currently overlap with the forests surveyed and has been excluded from this study. We are primarily focused on forest damage caused by deer species, with a special attention on the characteristics that attract deer. We also are interested in how the relative abundances of the three deer species, either occurring as a single species or in interactions with co-occurring ones, contribute to the likelihood of forest damage. To disentangle the complex relationship between deer, forests, and damage, we collated a unique database comprising: (i) data on forest characteristics and deer damage (presence and type) from the National Forest Inventory (NFI), and (ii) data on deer relative abundance gathered by a recent nationally coordinated project (SMARTDEER, Morera-Pujol et al., 2023) which we describe below in full detail along with our data analysis and modelling approach.

### National Forest Inventory Data (NFI): Forest Characteristics and Deer Damage

We gathered forest characteristics and deer damage data from the NFI carried out by the Forest Service of the Department of Agriculture, Food, and the Marine between 2015 and 2017 across the entire nation (DAFM, 2017). The NFI’s main objective is to monitor the composition, health and change in forest cover for both private and public forests across Ireland (DAFM, 2017). A permanent randomised systemic 2km x 2km grid that covered the entire land area of the Republic of Ireland created initial plot locations within the grid intersections (DAFM, 2017). The sampled plots (each circular plot measuring 25.24m in diameter, n = 1,681), which are a subset of all available grid cells, were then randomly assigned within 100m of each intersection and classified into land-use types to identify forest locations to be surveyed (DAFM, 2017) (Fig. 1). Forest characteristics accessible from NFI databases detail the biophysical attributes of the plot’s location. In more detail, the covariates extracted from the NFI were: altitude (metres); development stage (eight levels ranging from recently established forest to mature stands); European forest type (EFT: broadleaf, conifer, mixed); establishment type (afforestation, reforestation, semi-natural); forest availability for wood supply, which describes if a forest is in a suitable condition for harvesting wood (FAWS: available, unlikely, and not available); forest subtype (pure and mixed); canopy cover (%); mixture type (uniform, individually mixed, group mixed); nativeness (native, mixed, non-native); soil group (11 different soil groups); tree species (104 recorded species); stocking number (number of trees present in the sampling area); thin status (frequency and history of harvesting broken down into seven categories); tree distribution (regular, group, random); woodland habitat (semi-natural woodland, highly modified/non-native woodland, scrub/transitional woodland); slope (°); number of tree layers (1, 2, multistoried); bryophyte cover (%); fern cover (%); grass cover (%); herb cover (%); shrub cover (%); vegetation cover (%); and the age of each tree recorded. Furthermore, the NFI recorded the presence or absence of deer damage (see Fig 1 for the spatial distribution across Ireland). Damage instances were further classified into three categories: browsing (consumption of plant and tree shoots and leaves), bark stripping (peeled or torn stem and bark with broad teeth marks running down the tree), and tree fraying (male deer rubbing or fraying the velvet off their antlers and/or mark their territory). We computed a binary variable for overall damage by coding any type of damage as 1 and absence of damage as 0; we also coded individual binary variables for the three specific types of damage (presence of that particular type of damage = 1, absence of that particular type of damage = 0, with types of damage being either browsing, bark stripping, or fraying). Data collection has been carried out by expert foresters who have received extensive training and have long-term experience with vegetation data collection, reducing the likelihood of, for example browsing, to be underreported to levels that we consider negligible in our modelling effort. The NFI data were validated throughout the fieldwork period with 98 plots randomly surveyed between April 2016 and June 2017 (DAFM, 2017). Fieldwork validation scored a “good” (highest score) rating at 64% representing high quality data collection (DAFM, 2017). Only 7% of sites re-surveyed were classified as “re-measure” (DAFM, 2017).

**Fig. 1:**
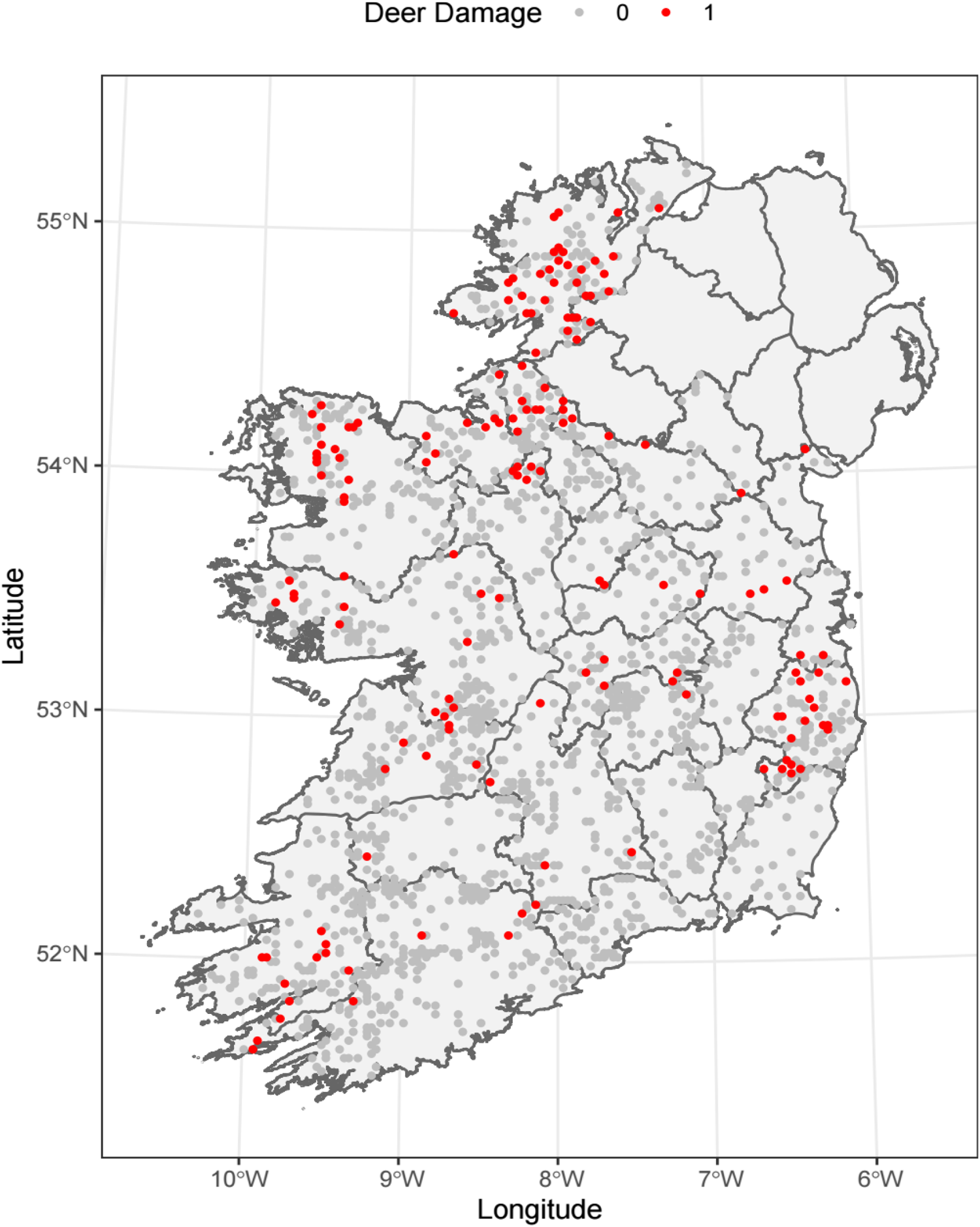
Map of Ireland showing the distribution of plots monitored by the 2017 National Forest Inventory (NFI) classified by presence (1) or absence (0) of deer damage to forests. Deer damages are further classified as bark stripping, browsing, and fraying, see methods for full details.

### SMARTDEER Project Data: Relative Abundance of the Three Species of Deer

Morera-Pujol et al., (2023) collated data on deer species for the last decade in Ireland, and, using recent advances with Bayesian integrated species distribution models, produced nationwide relative distribution maps for red, sika and fallow deer (Fig 2). These relative distributions (scaled to range between 0 and 1) were combined with National Parks and Wildlife Service (NPWS) culling return data (e.g., number of deer harvested nationwide for each species) to have, at the 5 x 5 km pixel level, the relative abundance of deer species (Morera-Pujol et al., 2023): as a result, for a given pixel, for instance, 1 red deer can be predicted to occur with 0 fallow deer and 15 sika deer. While the SMARTDEER original models contained uncertainties, which we could not incorporate into our own models, they were of high quality and accuracy and validated with independent deer data (Morera-Pujol et al., 2023). We used these up-to-date and comprehensive spatially explicit deer data as important predictors in our models explaining the variability of deer damage. We acknowledge that there is temporal variation between the two datasets (NFI data: 2015-2017, SMARTDEER: 2012-2022) used in this study, however these are the two most up to date datasets that are available regarding forest inventory and deer distributions. We later discuss and stress the need for further cooperation among stakeholders and government agencies, and how this would help reduce such temporal differences and enhance modelling reliability.

**Fig. 2:**
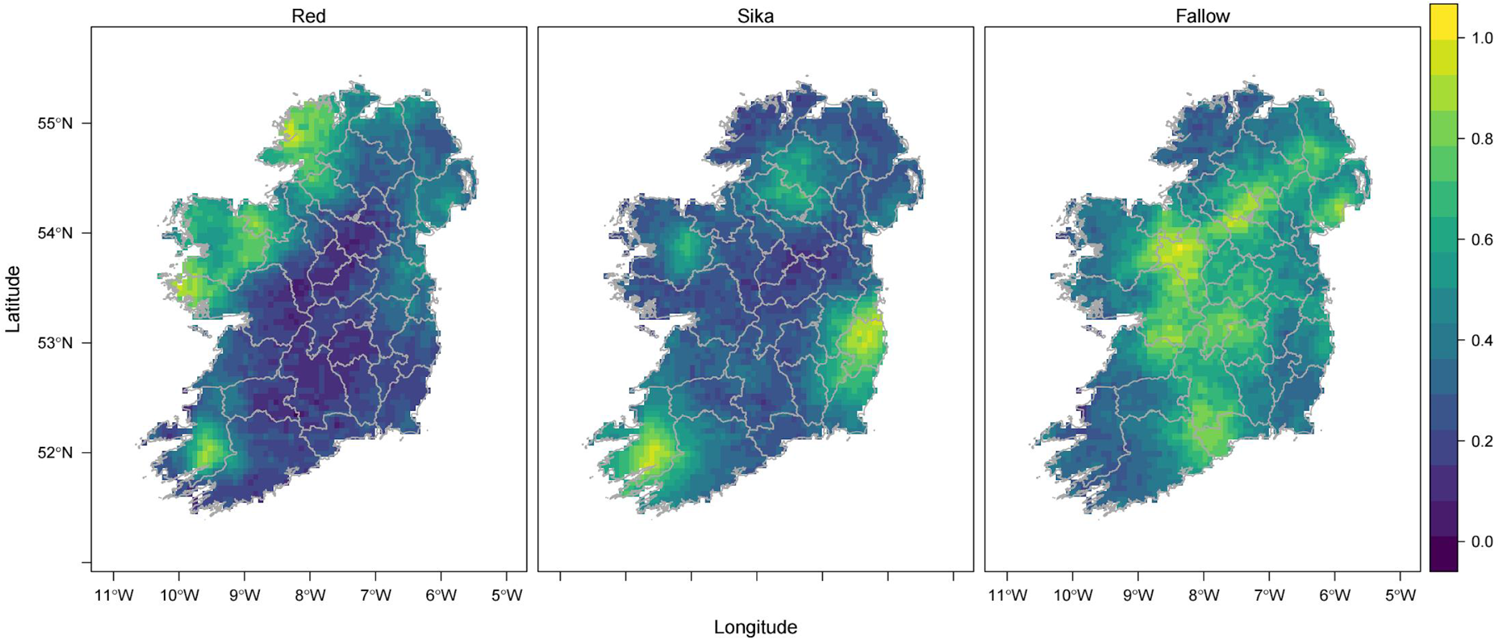
Map of Ireland showing the distribution and relative abundance for: a) red deer, b) sika deer, and c) fallow deer as predicted by Morera-Pujol et al., (2023). The values of the colour gradient represent relative abundance of deer at the pixel level (5 x 5km), rescaled to range from 0 to 1 to improve readability. The value 1 corresponds to the highest relative abundance for a given species recorded in a given pixel across Ireland. The three species are on the same scale, meaning for example that sika deer is predicted to be two times more abundant than red deer when the two species are assigned with relative abundance 0.6 and 0.3, respectively.

### Data Analysis

Data handling and analyses were carried out in R 4.3.0 (R Core Team, 2021). Ecological variables were computed based on the initial raw data from the NFI listed above. To capture the diversity of tree species, which is a good indicator on forest type (i.e., very low richness likely a single tree species plantation) and overall ecosystem health, richness (number of the different tree species) and Shannon diversity index were calculated using the package ‘vegan’ (Oksanen et al., 2020). To capture the arboreal cohort of each surveyed plot, average age (proxy for overall age of the forest stand, i.e., young vs mature) and standard deviation (highlighting the heterogeneity of the forest stand, i.e., young trees vs old trees vs heterogeneous) were calculated from the values of age of each single tree recorded in the forest plots. Prior to running any models, all candidate variables expected to explain the variability in deer damage across Ireland were screened for collinearity (with a threshold for the Pearson correlation coefficient set to |*rp*| < 0.5) (Dormann et al., 2013). Species richness was collinear with Shannon index and canopy cover, so we retained species richness. Vegetation cover (%) was collinear with bryophyte, grass, and shrub cover (%) and, to keep a variety of variables detailing the floral composition of each plot within the model, we retained bryophyte, grass, and shrub cover (%). EFT (European Forest Type) was collinear with woodland habitat, and we retained EFT. Nativeness was collinear with tree distribution, establishment type and FAWS and we retained nativeness. Mixture type was collinear with the forest subtype, and we retained mixture type. To fully account for likely spatial correlation in the data, latitude, longitude, and their interaction were included as predictors in all the models. Finally, the variable for number of layers was removed from analysis. Number of layers refers to the number of layers within the canopy with the first two layers recorded as “1” or “2” and anything above “2” was recorded as “multistoried”. This unbalance caused disruption to the model performance and was therefore removed at a later stage of analysis.

Since both average age and its standard deviation provide relevant information on the age and structure of forests, but were collinear, we performed a Principal Component Analysis (PCA) to include them both in the model (Fig 3). The two resulting uncorrelated principal components, PC1 and PC2, explained 70 and 30% of the total variance respectively, and were both included in the model structures. PC1 increases with decreasing average age and standard deviation (SD) of age, and thus classifies the forest either as young plantation (low average age & low SD age) or mature natural woodland (high average age & high SD age). PC2 increases with decreasing average age and increasing SD, thus classifying the forests in a spectrum from mature even-aged plantation (high average age & low SD age) to young natural woodland (low average age & high SD age). Finally, PC1 was also collinear with the development stage, and we retained PC1. The final list of covariates selected after the collinearity screening were altitude (m), slope (°), bryophyte cover (%), fern cover (%), grass cover (%), herb cover (%), shrub cover (%), EFT, stocking, mixture type, nativeness, species richness, PC1, PC2, the relative abundance of red deer, sika deer and fallow deer, and latitude and longitude.

**Fig. 3:**
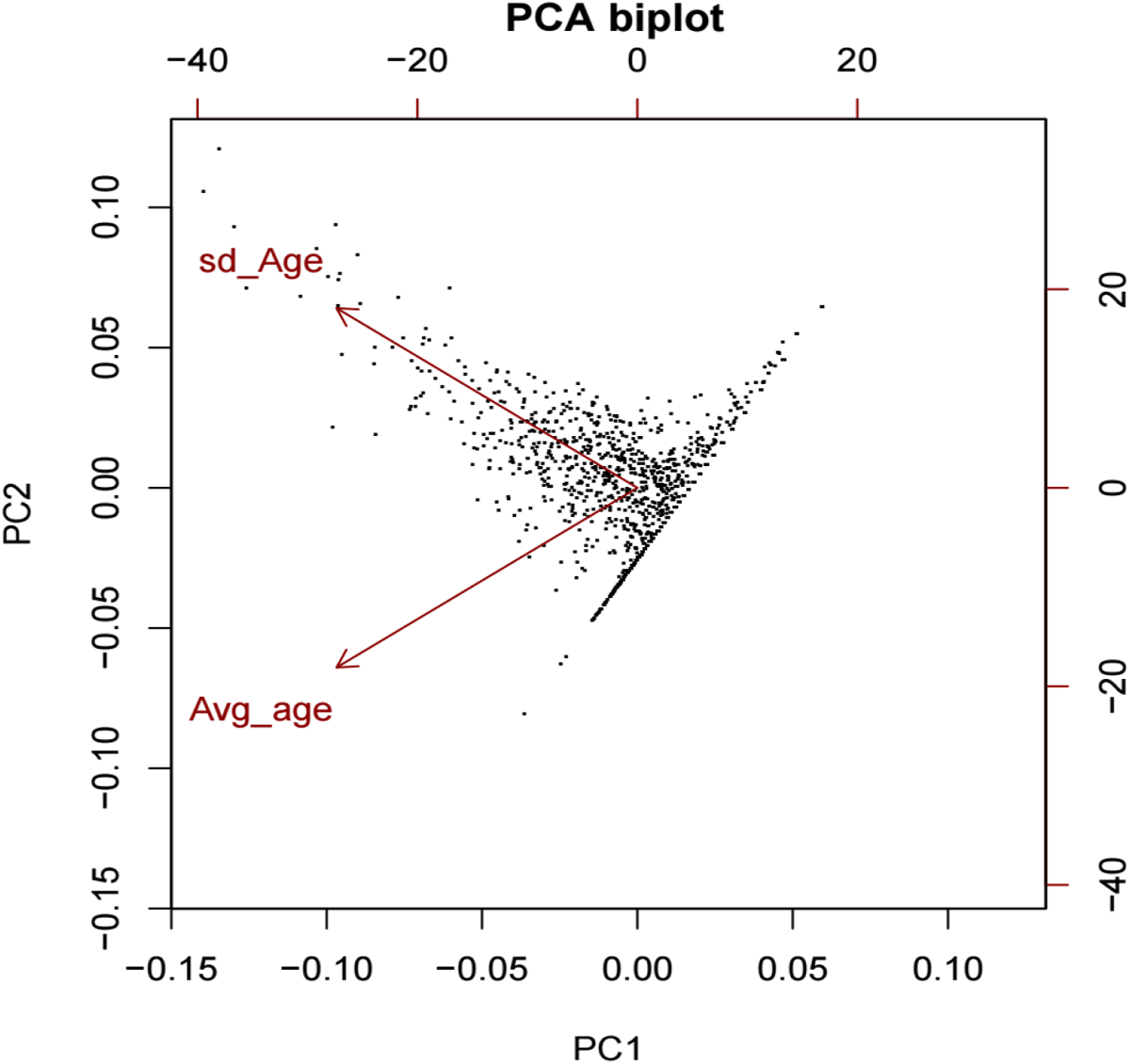
Principal Component Analysis (PCA) biplot embracing the variability of average and the standard deviation of tree age for each forest plot. As PC1 increases from negative to positive values the characteristic of the forest changes from a mature natural woodland (older forest with high variability in age) to a young plantation (younger forest with no variability in age). As PC2 increases from a negative value to a positive value the characteristics of the forest changes from a mature plantation (older forest with no variability in age) to a young natural woodland (young forest with high variability in age).

We ran four binomial generalized linear models (GLMs) with presence/absence of damage as a response (overall deer damage; bark stripping damage; browsing damage; fraying damage) and with the relative abundance of native and non-native deer species and environmental and forest characteristics listed above as covariates. All numeric predictors were scaled and included as quadratic terms to allow for non-linear effects. The relative abundance of each deer species was included as a single (and quadratic) term and also as an interaction between each species (i.e., red interacting with sika, red interacting with fallow and sika interacting with fallow) to capture potential interaction effects. During preliminary analyses, the models including the full structure performed well and had no convergence issues. To present a more parsimonious model structure, however, we applied the *step* function (Hastie and Pregibon, 1992; Venables and Ripley, 2002) which simplified the model structure by excluding those variables that did cause an increase in the Akaike Information Criteria (AIC) value when included in the model. The absence of spatial autocorrelation was confirmed with a variogram of the residuals (Fig S1, S2 and S3 for the 3 GLMs, respectively) (Pebesma, 2004; Pebesma and Bivand, 2005). We used the DHARMa package to ensure our models do not violate any of the model assumptions (Fig S4, S5, S6 for the 3 GLMs, respectively) (Hartig, 2022). The predictions of the models were plotted using the ‘effect’ and ‘jtools’ library (95% marginal confidence interval, CI) (Cohen et al., 2003; Bauer and Curran, 2005; Fox, 2003; Fox and Weisberg, 2018; 2019). Using the model predictions for the different type of damage, we created risk scenarios depicting the likelihood of damage as a function of clear effects (95% CI not overlapping zero) of the deer species relative abundances and corresponding interactions. Finally, the model predictions were also used to create predictive damage maps for each surveyed plot across the nation.

## Results

### Modelling Deer Damage

Of the four models run, the initial model for overall deer damage performed the worst in comparison with the other three single damage type models based on the AIC (AIC = 885; ΔAIC between this model and the other three is greater than 300, see Table 1). This result confirms the importance of recording the specific damage type, as opposed to overall damage, in forest surveys. Nevertheless, the model for overall deer damage showed clear effects in several variables indicating damage was more common towards northern latitudes and in areas with more bryophytes, grass, trees, higher tree richness, lower PC1 values (i.e., more natural woodlands) and greater sika deer likelihood (Table 1). Overall deer damage was also found to be more common in areas with less herbs and shrubs (Table 1). Finally, overall damage was also significantly driven by the interactions among the three different deer species (Table 1). However, because of the poor performance of this model compared to those tackling the different types of deer damage individually, no further inference is drawn from this starting model, and the rest of this result section will focus on the single damage type models.

**Table 1.** Generalized linear models predicting the likelihood of overall deer damage, bark stripping, browsing, and fraying across 1,681 sampling stations within Irish forests from 2015 to 2017. Odds ratios (OR) were obtained by exponentiating model coefficients in the linear scale, with values under 1 representing negative effects of the covariate, whereas values over one positive effects; standard error (SD), 95% confidence intervals (CI), p-values (p), Akaike information criterion (AIC) and proportion of variance explained (R^2^) were also reported in full.

|  | Overall Deer Damage |  |  |  | Bark Stripping Damage |  |  |  | Browsing Damage |  |  |  | Fraying Damage |  |  |  |
| --- | --- | --- | --- | --- | --- | --- | --- | --- | --- | --- | --- | --- | --- | --- | --- | --- |
| Predictors | OR | SD | CI | p | OR | SD | CI | p | OR | SD | CI | p | OR | SD | CI | p |
| Intercept | 0 | 0 | 0.00 – 0.00 | <0.001 | 0 | 0 | 0.00 – 0.00 | <0.001 | 0 | 0 | 0.00 – 0.01 | <0.001 | 0 | 0 | 0.00 – 0.00 | 0.009 |
| Latitude (km) | 1.01 | 0 | 1.01 – 1.01 | <0.001 | 1.01 | 0 | 1.00 – 1.01 | <0.001 | 1.01 | 0 | 1.01 – 1.02 | <0.001 | 1.07 | 0.03 | 1.02 – 1.14 | 0.016 |
| Longitude (km) |  |  |  |  |  |  |  |  | 0.99 | 0 | 0.98 – 1.00 | 0.066 | 1.09 | 0.04 | 1.02 – 1.18 | 0.026 |
| Lat:Long (km) Interaction |  |  |  |  |  |  |  |  |  |  |  |  | 1 | 0 | 1.00 – 1.00 | 0.025 |
| Altitude (km) | 0.82 | 0.11 | 0.64 – 1.05 | 0.127 |  |  |  |  |  |  |  |  | 0.65 | 0.15 | 0.40 – 1.02 | 0.07 |
| Slope° |  |  |  |  | 0.76 | 0.13 | 0.51 – 1.02 | 0.113 |  |  |  |  |  |  |  |  |
| Tree Richness | 2.29 | 0.76 | 1.21 – 4.48 | 0.013 | 2.74 | 1.23 | 1.17 – 6.87 | 0.025 | 5.16 | 2.99 | 1.76 – 17.38 | 0.005 |  |  |  |  |
| Tree Richness<sup>2</sup> | 0.62 | 0.19 | 0.33 – 1.09 | 0.109 | 0.52 | 0.22 | 0.21 – 1.14 | 0.127 | 0.39 | 0.19 | 0.13 – 0.96 | 0.056 |  |  |  |  |
| Tree Stocking | 1.6 | 0.17 | 1.30 – 1.97 | <0.001 |  |  |  |  | 1.37 | 0.21 | 1.02 – 1.86 | 0.039 | 1.95 | 0.39 | 1.33 – 2.89 | 0.001 |
| Tree Stocking<sup>2</sup> |  |  |  |  | 0.08 | 0.09 | 0.01 – 0.66 | 0.025 |  |  |  |  |  |  |  |  |
| PC1 | 0.79 | 0.09 | 0.63 – 0.99 | 0.039 | 0.72 | 0.13 | 0.51 – 1.03 | 0.068 |  |  |  |  |  |  |  |  |
| PC1<sup>2</sup> |  |  |  |  |  |  |  |  |  |  |  |  | 0.19 | 0.16 | 0.03 – 0.70 | 0.045 |
| PC2 | 0.84 | 0.1 | 0.67 – 1.06 | 0.151 | 0.56 | 0.1 | 0.39 – 0.79 | 0.001 | 1.29 | 0.22 | 0.91 – 1.79 | 0.148 |  |  |  |  |
| Bryophyte Cover % | 1.56 | 0.2 | 1.22 – 2.03 | 0.001 | 1.49 | 0.27 | 1.06 – 2.15 | 0.025 |  |  |  |  | 0.28 | 0.21 | 0.07 – 1.38 | 0.083 |
| Bryophyte Cover<sup>2</sup> % |  |  |  |  |  |  |  |  |  |  |  |  | 5.69 | 3.97 | 1.36 – 21.95 | 0.013 |
| Fern Cover<sup>2</sup> % |  |  |  |  |  |  |  |  | 1.45 | 0.23 | 1.06 – 1.98 | 0.018 |  |  |  |  |
| Grass Cover % |  |  |  |  |  |  |  |  | 2.26 | 0.47 | 1.52 – 3.47 | <0.001 | 0.22 | 0.14 | 0.06 – 0.78 | 0.02 |
| Grass Cover<sup>2</sup> % | 1.35 | 0.15 | 1.09 – 1.67 | 0.006 |  |  |  |  |  |  |  |  | 5.34 | 3.48 | 1.51 – 19.69 | 0.01 |
| Herb Cover % | 1.82 | 0.57 | 0.99 – 3.37 | 0.056 | 2.82 | 1.12 | 1.32 – 6.26 | 0.009 |  |  |  |  | 2.46 | 1.55 | 0.73 – 8.77 | 0.156 |
| Herb Cover<sup>2</sup> % | 0.49 | 0.16 | 0.26 – 0.91 | 0.027 | 0.35 | 0.15 | 0.15 – 0.77 | 0.012 |  |  |  |  | 0.3 | 0.21 | 0.07 – 1.10 | 0.086 |
| Shrub Cover % |  |  |  |  | 2.17 | 1.07 | 0.83 – 5.85 | 0.119 |  |  |  |  |  |  |  |  |
| Shrub Cover<sup>2</sup> % | 0.72 | 0.08 | 0.57 – 0.90 | 0.005 | 0.32 | 0.16 | 0.12 – 0.83 | 0.021 |  |  |  |  |  |  |  |  |
| Red Deer Relative Abundance | 1.15 | 0.15 | 0.89 – 1.49 | 0.283 | 1.79 | 0.33 | 1.25 – 2.57 | 0.002 | 20.3 | 34.72 | 0.84 – 699.84 | 0.078 | 0.91 | 0.29 | 0.47 – 1.70 | 0.762 |
| Red Deer Relative Abundance<sup>2</sup> |  |  |  |  |  |  |  |  | 0.04 | 0.06 | 0.00 – 0.75 | 0.043 |  |  |  |  |
| Fallow Deer Relative Abundance | 1.04 | 0.13 | 0.81 – 1.32 | 0.75 | 6.56 | 6.93 | 0.90 – 57.04 | 0.075 | 1.36 | 0.3 | 0.88 – 2.12 | 0.163 | 1.32 | 0.32 | 0.81 – 2.13 | 0.254 |
| Fallow Deer Relative Abundance<sup>2</sup> |  |  |  |  | 1.58 | 0.2 | 1.23 – 2.02 | <0.001 |  |  |  |  |  |  |  |  |
| Sika Deer Relative Abundance | 1.57 | 0.21 | 1.21 – 2.02 | 0.001 | 1.06 | 0.23 | 0.68 – 1.59 | 0.777 | 0.32 | 0.31 | 0.05 – 2.08 | 0.232 | 1.89 | 0.45 | 1.17 – 3.00 | 0.008 |
| Sika Deer Relative Abundance<sup>2</sup> |  |  |  |  |  |  |  |  | 5.94 | 5.4 | 0.99 – 35.58 | 0.05 |  |  |  |  |
| Red-Sika Interaction | 0.68 | 0.06 | 0.57 – 0.81 | <0.001 | 0.75 | 0.09 | 0.58 – 0.94 | 0.015 | 0.63 | 0.1 | 0.44 – 0.86 | 0.006 | 0.7 | 0.16 | 0.44 – 1.07 | 0.115 |
| Red-Fallow Interaction | 0.73 | 0.09 | 0.57 – 0.92 | 0.01 | 1.4 | 0.27 | 0.96 – 2.04 | 0.082 | 0.37 | 0.11 | 0.20 – 0.65 | 0.001 | 0.43 | 0.11 | 0.26 – 0.69 | 0.001 |
| Sika-Fallow Interaction | 0.58 | 0.09 | 0.43 – 0.77 | <0.001 | 0.51 | 0.13 | 0.30 – 0.81 | 0.007 | 0.43 | 0.11 | 0.25 – 0.69 | 0.001 |  |  |  |  |
| $R^2$ | | | 23.73 % | | | | 28.32 % | | | | 37.6 % | | | | 26.94 % | |
| AIC |  |  | 885 |  |  |  | 574 |  |  |  | 366 |  |  |  | 340 |  |

Compared to the overall deer damage model, the bark stripping model was characterized by a significantly lower AIC score (AIC = 574), and variables such as grass cover and altitude were dropped by the model simplification process, whereas slope was this time included (Table 1). Bark stripping damage was more common towards northern latitudes, in areas with decreasing PC2 values (i.e., mature plantation; Table 1, effect plot in Fig. S7), and in areas with more abundant bryophytes, trees, tree richness and red deer (for single deer species effect plots see Fig. S10). There was a negative quadratic relationship between bark stripping damage and herb and shrub cover highlighting higher bark stripping damage in areas characterized by intermediate values of these variables (Table 1, effect plots in Fig. S7), as well as fallow deer relative abundance (for single deer species effect plots see Fig. S10). Bark stripping damage was also driven by the interaction between red and sika deer, and sika and fallow deer (interaction effects among deer species will be discussed below in the ‘Risk scenarios’).

Based on another significant drop in AIC score (AIC = 366), the browsing model performed better in comparison to the bark stripping model. Variables such as PC1 and bryophyte, herb and shrub cover were dropped by the model simplification process (Table 1). New variables including longitude, fern and grass cover were this time included. Browsing damage was more common in areas with higher tree richness, grass cover, ferns cover, trees (Table 1, see effects plots in Fig. S8) and sika deer (for single deer species effect plots see Fig. S10). Browsing damage was also driven by the interaction of all three deer species (interaction effects among deer species will be discussed below in the ‘Risk scenarios’).

The model for fraying damage was the one with the lowest AIC score (AIC = 340). Compared to the previous models, variables dropped included tree species richness, PC2, fern and shrub cover as well as the interaction between sika and fallow deer, whereas the model simplification process retained PC1 and bryophyte cover (Table 1). PC1 had a negative quadratic effect on fraying damage (Table 1, Fig S9), showing that damage is less likely at both mature natural woodlands and young plantations, while slightly more common in mixed forests with a mid-age range. Slightly higher bryophyte and grass cover at sites were linked to higher fraying damage (Fig S9). Higher tree stocking was also linked to greater fraying damage (Fig S9). Finally, higher Sika deer relative abundances were linked to higher fraying likelihood. The interaction between red and fallow deer also affected the likelihood of fraying damage across the country (interaction effects among deer species will be discussed below in the ‘Risk scenarios’).

### Risk Scenarios

For each deer damage model, we created risk scenarios depicting the likelihood of damage at different relative densities of multiple co-occurring deer species. To select the scenarios, we considered only deer species with a significant effect (95% CI not overlapping zero) for their interaction in the corresponding model (Table 1 – for interaction effect plots see Fig S11). The most relevant risk scenarios are reported in Fig. 4. Note that the 1,681 sampling stations spread across Ireland span along the data space of each deer species (meaning that at least one sampling station is present across the different levels of relative deer abundance). The number of sampling stations available at extreme deer densities, however, is generally lower than that available for intermediate densities, therefore deer damage predictions must be taken under careful consideration wherever the models have been trained with reduced sample sizes. Our scenarios in Fig. 4 reflect this conservative approach (less certain damage likelihood are reported by numbers in smaller font size) which implies avoiding over-interpreting data (and wrongly informing stakeholders and policy makers). Moreover, in most scenarios presented in Fig 4, when a given deer species is occurring at greater relative abundance, it is less likely to have data from sampling stations where another species is also occurring at increasingly higher density. This confirms what shown in Fig 2 with clear distinctive and non-overlapping hotspots of presence for the three deer species present in Ireland. All species, when occurring a relatively lower abundances, are responsible for minor damage to forest. Our models however predict quite complex scenarios and heavy browsing damage when the highest sika deer densities are recorded (Fig. 4). Likewise, fraying damage is linked to higher fallow deer densities, bark stripping damages to higher sika and red deer densities.

**Fig. 4:**
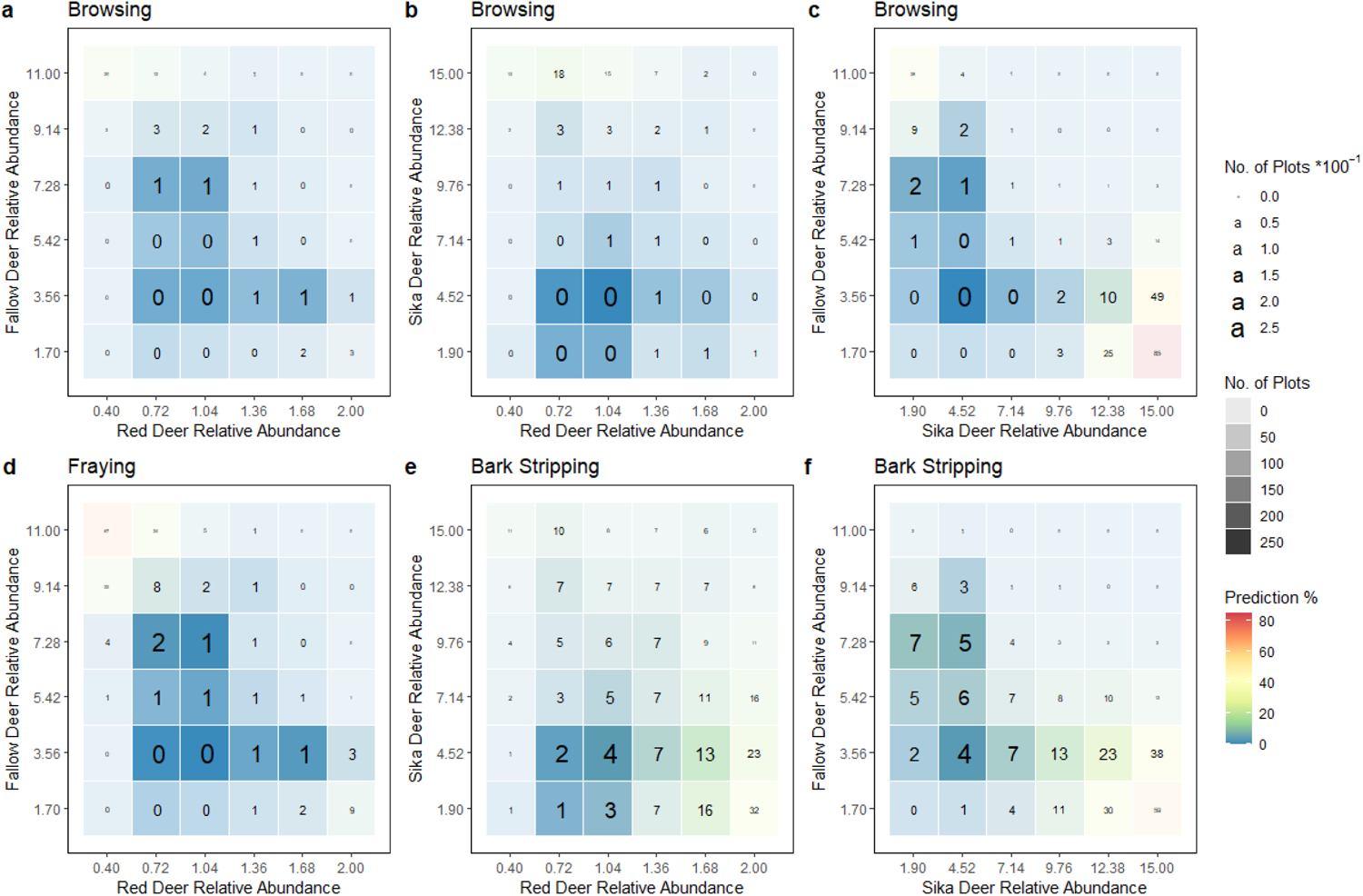
Risk scenarios predicted by the generalized linear models at interactive relative abundances for: a) browsing damage for red and fallow deer; b) browsing damage for red and sika deer; c) browsing damage for sika and fallow deer; d) fraying damage for red and fallow deer; e) bark stripping damage for red and sika deer; and f) bark stripping damage for sika and fallow deer. Numbers inside plots refer to predicted damage risk (in percentage: cold blue/green colours indicating less damage than warm orange/red colours) whereas the size of the text as well as the level of background shading are proportional to the sample size used to train the model (i.e., reduced confidence on the predicted scenarios when sampling stations are lower in sample size).

Finally, based on the predictions from each of the single damage models, we created predictive damage maps by forecasting the likelihood of each damage type at all the sampled locations across Ireland (Fig. 5). A high likelihood of damage was predicted for bark stripping (Fig. 5a) in hotspot areas in the northwest, southwest, and east of the nation (which are all either sika or red deer hotspots, respectively, see Fig 2 with distribution of deer hotspots for a visual comparison), whereas we predicted lower damage likelihood across the midlands, where fallow deer is the dominant species (Fig. 5a). Predicted likelihood of browsing damage was quite similar to bark stripping with more intense damage found in parts of the midlands (Fig 5b). Lastly, the predictions for fraying again showed higher likelihood of damage in the hotspot areas in the northwest and east of the country, but very low likelihood in the southwest (Fig 5c).

**Fig. 5:**
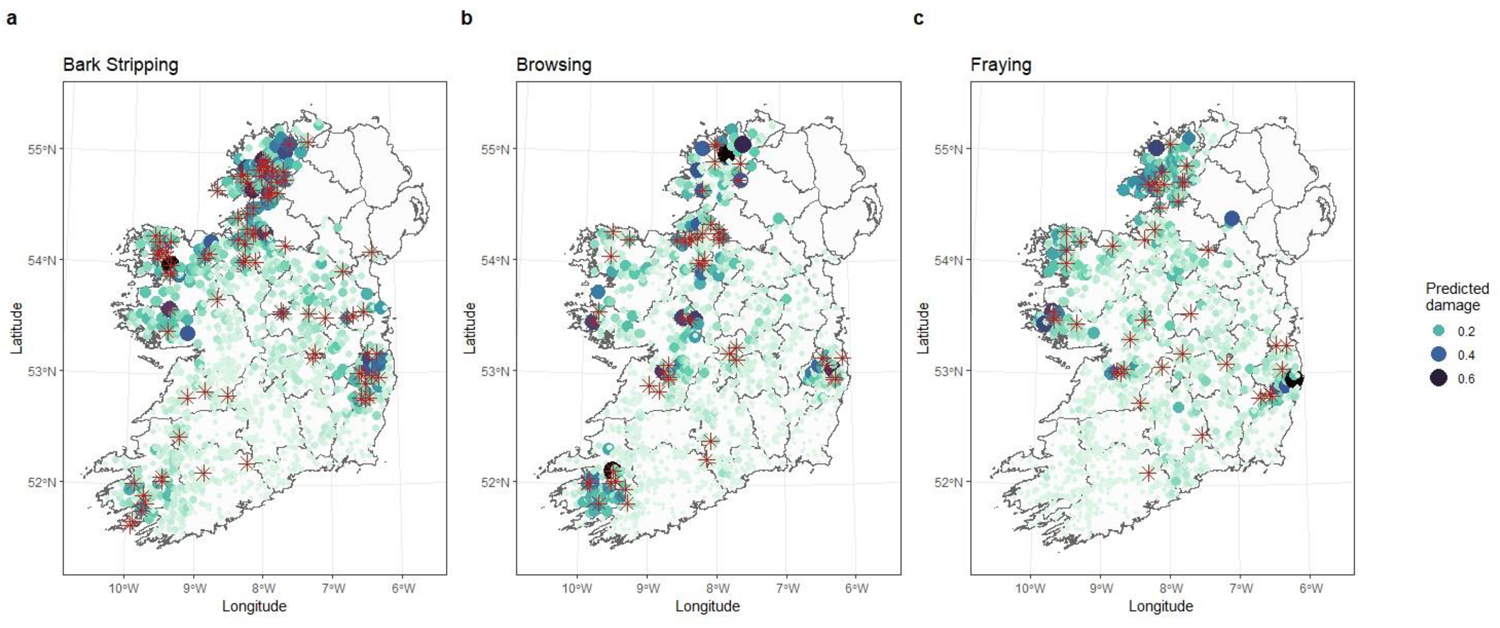
Predicted percentage damage for the 1,681 sampling stations across Ireland, based on the outcome of the generalized linear models, for: a) bark stripping damage, b) browsing damage, and c) fraying damage. Locations where damage was observed are indicated by the red star (*) for each respective damage type. Both symbol size and colour gradients represent the predicted damage likelihood, with smaller and lighter green/blue colours indicating lower levels of damage and larger, darker colours representing higher levels of damage.

## Discussion

Our study combined multiple complimentary data sources across a national landscape such as integrated deer species distribution models (Morera-Pujol et al., 2023) and forest inventory data to investigate the role of multiple (both native and non-native) deer species on the presence, type, and severity of forest damage. We deduced the role of forest characteristics, deer species and ecological interactions that drive three types of damage (bark stripping, browsing, and fraying) in forests ecosystems. Forest characteristics played a key role in the severity and type of damage risk that deer posed. Dense forests are the ideal forests for all three types of damage. In mature forests, with high tree diversity and ground level flora (e.g., bryophytes, herbs, and shrubs) we found that bark stripping was the primary form of damage. Browsing damage was more common in areas with increased grass and fern cover with higher tree diversity. In immature mixed woodlands we found that fraying was more common, however, the importance of ground level flora (e.g., bryophytes and grass) was also supported by our models.

Across all our study sites we found that forest damage (type, severity) was shaped by interactions of multiple deer species occurring simultaneously, suggesting subtle inter-species competition and exclusion dynamics also observed in other ungulate populations that require further investigation to understand the ecological mechanism (Zini et al., 2023). After accounting for environmental and forest characteristics that attract deer species and increase the likelihood of damage, we disentangled which species are more responsible for causing forest damage and obtained insights into why damage occurred. We did this by inspecting the interactions among deer species and the damage dynamics occurring when more than one species was present. We used our modelling approach to provide simulated risk scenarios that offer pragmatic guidance for forest managers to reduce the risk of extensive damage from deer on their property. We provide a framework for long term and large-scale monitoring of the impacts that multiple deer species have on the national forest estate and this approach is transferable to any initiative that aims to monitor forest damage by multiple sympatric deer species outside Ireland.

### Ecological Drivers of Damage

Our analysis demonstrates that across national heterogenous forests there are specific characteristics that appear more attractive to deer and thus, place these forests at higher risk for species specific behaviours that cause considerable damage. We found that dense woodlands with understory vegetation such as high bryophyte cover were at a higher likelihood of both bark stripping and fraying damage occurring. Bryophytes are small, mossy, non-vascular plants that are sensitive to habitat change, such as that related to intensive forest plantations (Scheidegger & Werth, 2009) and their abundance generally increases alongside stand age, up to 30 years, which may suggest that deer species known for bark stripping and fraying tree stands may prefer more established woodlands (Schmalholz & Hylander, 2009; Morera-Pujol et al., 2023; Murphy et al., 2023). Furthermore, deer species selection of dense forest habitats is driven by the need to find refuge under heightened hunting pressure, further emphasizing their preference of dense forest systems (Lone et al., 2015; Murphy et al., 2023). Our analysis certainly points to this phenomenon as we found that bark stripping and fraying damage is dependent on age structure: bark stripping in mature plantations, fraying in middle aged mixed woodlands highlighting a potential habitat preference for these behaviours. In contrast browsing was more common in more diverse woodlands where open areas that are covered in grass and ferns might be more common. Diverse forests provide deer with abundant resources for browsing and are more likely to have a larger proportion of preferential broadleaf trees (Gill, 1992) while mature trees in monoculture plantations seem more suited for bark stripping behaviour. The data used in our modelling effort were collected at the site level, each plot including up to a maximum of 512 single trees, but the species of the tree where the actual deer damage was observed was not available. Our models could be much more powerful with information on the species and age of the actual tree damaged, helping managers with prediction at a greater resolution. We suggest that these data should be collected and analysed in future inventory schemes.

### Single and Sympatric Deer Species

We found that high deer relative abundance increases the likelihood of damage in forests, particularly when more than one species co-occurs. Deer species have previously been shown to cause damage to forests either as a single species or within a multi-species system (e.g., Chianucci et al., 2015; Welch and Scott, 2017; Zini et al., 2022). Our study provides strong support to those results by introducing the use of accurate relative abundance estimates for three co-occurring deer species as predictors in the models. First, relative abundance of non-native sika deer was the main driver in the likelihood of browsing and fraying damage. Sika deer are opportunistic foragers and, in combination with their ability to successfully interbreed with native red deer populations, have proliferated in established hotspot areas across the country (McDevitt et al., 2009; Purser et al., 2010; Morera-Pujol et al., 2023). Sika deer are primarily found in the east and southwest of Ireland, with smaller populations also present in the red deer hotspots of the northwest, coinciding with historical introductions (Powerscourt 1884; Carden et al., 2011; Morera-Pujol et al., 2023). Forests in such areas dominated by overabundant deer (sika occurring 7-fold higher than red deer) are likely to experience high levels of damage. The impacts of overabundant sika populations alone can cause severe ecological damage such as aiding the expansion of invasive species, (e.g., *Rhododendron,* Yokota et al., 2009), increase mortality of tree seedlings (Tsujino and Yumoto, 2004), and alter the composition of insect and soil communities (Akaba et al., 2014; Tsurumi et al., 2015). Management of sika and red deer populations throughout the country should be of the highest importance to mitigate further ecological damage.

Secondly, the effect of all three deer species in forests is exacerbated when more than one species co-occurs. The general pattern found among the interactions between species shows damage is likely when the relative abundance of a single species is high. When the relative abundance of both sympatric species is moderate the likelihood of damage is relatively low but increases dramatically with small increases to one of the other species’ relative abundances. This is the case of, e.g., browsing by fallow and sika deer, two non-native species. The highest predicted likelihood of damage is the highest across all scenarios but, at low and moderate levels of these species’ relative abundance, damage is extremely limited. These low and moderate abundance scenarios are common throughout Ireland. Damage dramatically increases with small increases in relative abundance of either fallow or sika (not both, since the two species only meet at the edges of their respective distributions) as ecological heterogeneity is lost. Such scenarios may be less common but should be given highest priority for managing deer populations across the country, particularly given the recent explosion of deer populations throughout the country (Carden et al., 2011, Morera-Pujol et al., 2023, Murphy et al., 2023) that may lead to these scenarios becoming more widespread if the required mitigation measures are not implemented accordingly.

Species that have co-evolved with one another typically partition resources to avoid competition (e.g., Pianka, 1973; Jenkins and Wright, 1988; Redjadj et al., 2014). To avoid competition, two species may partition their resources in different ways (Stewart et al., 2010). Spatial partitioning may occur between red/sika deer (note distribution overlap, Fig. 2) and fallow deer for example. However, along with competition, spatial partitioning may also be due to species specific habitat preferences, which has been observed within ungulate species between sika deer and Japanese serows (*Capricornis crispus*), and within different *Cervidae* species such as mule deer (*Odocoileus hemionus*) and white-tailed deer (*Odocoileus virginianus*) (Seki and Hayama, 2021; Staudenmaier et al, 2021). Trophic partitioning may also occur where one species is more specialised in a particular feeding behaviour resulting in excessive damage that may force other species to find alternative resources. For example, at high densities in Japan, sika deer’s heavy browsing effects have been shown to deform forest structures and alter the composition of understorey vegetation, promoting the expansion of plants that are unpalatable for deer species (Shimoda et al., 1994; Takatsuki and Gorai, 1994). Similarly in Ireland, sika deer densities have been rapidly growing and their heavy browsing activity may lead to distorted forest structures that mirror the documented impacts observed in Japan. This may lead to the proliferation of plants that are unpalatable for other deer species, further distorting Ireland’s forest ecosystems. However, the three species of deer currently in Ireland have evolved separately from each other, and thus they may lack adaptations for coexistence (e.g., Putman, 1996; Forsyth and Hickling, 1998; Ferretti et al., 2011; Ferretti et al., 2015). Competition may arise between two or more species within the same trophic level when common resources are limited (de Boer and Prins, 1990), and when there is spatial and temporal variation of a common resource, especially within human-modified landscapes (e.g., Arsenault and Owen-Smith, 2002; Robertson et al., 2013), which affects the behaviour, growth, and survival of individuals (e.g., Durant, 1998; Harris and Siefferman, 2014). Our predicted likelihood of damage across the country shows high and intense likelihood of both bark stripping and fraying damage in deer hotspot areas in the northwest and east of the country, with bark stripping damage also highly likely in the southwest, primarily in either native red or non-native sika deer hotspots. Browsing damage is likely to be more evenly spread across the country with intense damage likelihood found in all hotspots. Here we have provided the tools to help predict the location and intensity of three types of forest damage by three different species of deer, for the first time.

At the site level, further studies will be required to disentangle the role of deer relative abundance on damage levels when sympatric deer species occur within forests. We show that the likelihood of damage decreases when two species occur at high densities in a forest, possibly due to interspecific competition between the two species. Specifically, in sika deer hotspots with a low relative abundance of red deer the likelihood of damage is high. Once red deer become established in an area with high levels of sika the likelihood of damage decreases. While we were able to detect this effect, we are unable to identify the fine scale drivers of this mechanism. It may be driven by the displacement of smaller sized sika deer by the larger bodied red deer within the forests, which has been observed in other habitats between *Cervidae* species (e.g., Ferretti et al., 2008). It would be interesting to shed light on this interaction, particularly between red and sika deer that have a history of hybridization (e.g., Bartos, 1981). We recommend several methods to disentangle the ecology of two sympatric species occurring in forest plantations where damage level is dependent on relative abundance. Camera trap surveys are a low cost and relatively low survey effort intensity method to passively monitor forest sites where two or more sympatric species are likely to occur (Smith et al., 2022). Camera trap surveys provide key population metrics, deliver insights into the behavioural ecology of the species and capture data on species presence, damage type and species interactions, data which can be used to disentangle our findings. Furthermore, practitioners without access to field sites may attempt to understand the mechanism we describe using an agent-based modelling simulation approach (Murphy et al., 2020). Agent based models parameterized with species specific information from the literature (population metrics, movement rates, activity) and complimented with high-frequency input data such as national forest inventory data and local geospatial data (landscape scale habitats, human footprint indices etc.) can provide a tool for practitioners to not only understand the underlying mechanisms we have shown but also to experiment with the efficacy of multiple management strategies prior to spending time and resources implementing them in the field (Murphy et al., 2020).

### Continued Monitoring and Management Recommendations

A coordinated approach in collating and analysing data on three species of deer has been the foundation of this study. Morera-Pujol et al. (2023) collated presence absence data from government bodies (e.g., Coillte) based on surveys of property managers and additional faecal pellet surveys, and presence only data from various sources such as published studies and citizen science programs (e.g., iNaturalist), and developed their own bespoke tools for recording deer presence (online survey and mobile app; Morera-Pujol et al., 2023). The result of Morera-Pujol et al., (2023) are national distributions for all prevalent deer species in Ireland that acts as a national baseline for future work on these species; however, higher resolution studies would benefit from more detailed monitoring at the local scale. The Department of Agriculture and the Marine (DAFM) have been successfully gathering data since 2006 highlighting the composition, state, and health of the national forest estate (DAFM, 2017). Combining these two comprehensive datasets has allowed us to highlight novel insights into the relationship between three widespread deer species and three specific types of forest damage. This approach can be reproduced in other countries with similar presence absence and presence only data to enhance the understanding of deer impacts on forest ecosystems.

While this work is promising, further long-term, coordinated data collection and cooperation (between government bodies, forest managers, private landowners, hunters, citizen scientists etc.) is essential to monitor the changes in deer populations’ size and range, and ensure these insights are used effectively and efficiently. Effective management of deer should be done through the lens of habitat management, taking not just the deer into consideration but the entire forest ecosystem (Reimoser and Gassow, 1996; Vospernick and Reimoser, 2008; Reimoser et al., 2009). Knowledge of deer distributions, population numbers and behaviour is needed for an integrated management approach that can maintain deer populations at sustainable levels, protect the viability of the forest industry and income within rural areas, and maintain a diverse and balanced ecosystem (Mayle, 1999; Nugent, 2012). We encourage the relevant authorities / practitioners to i) further enhance the coordinated approach in collecting information on deer and forest characteristics, and ii) use these predictions to help decision making and implement the relevant management options to mitigate and alleviate forest damage and further human wildlife conflicts, both on local and national scales. This study has provided baseline information by using an analytical approach to identify the main drivers of forest damage, identify which deer species is most likely to cause damage, and highlight areas most vulnerable to damage on a national scale, which will guide authorities in making science based and effective management decisions.

## Acknowledgements

We are thankful to all agencies and contributors that provided data and answered enquiries. We thank John Redmond, Luke Heffernan, and the entire forest service (DAFM) for the NFI data collection. We thank Tony Quinn and Andrew McCullagh (DAFM) for their continued support throughout the project. We thank Barry McMahon and Charles Harper for their contribution to the SMARTDEER project and their insights to this project. This work is a part of the nationally coordinated project ‘A smart and open-science approach to monitor and analyse deer populations in the Republic of Ireland and set the scene for evidence-based deer management’ funded by DAFM, grant no. 2019R417 (2019 Research Call Instruments (II–V)). Additional thanks to the School of Biology and Environmental Sciences, University College Dublin for their continued support and to the Higher Education Authority for financial support during Covid-19 funded-extensions. Finally, we thank the various foresters, farmers and hunters who gave valuable feedback during the presentation of this work to stakeholders.

