## Supplementary Materials for "Predicting forest damage using relative abundance of multiple deer species and national forest inventory data"

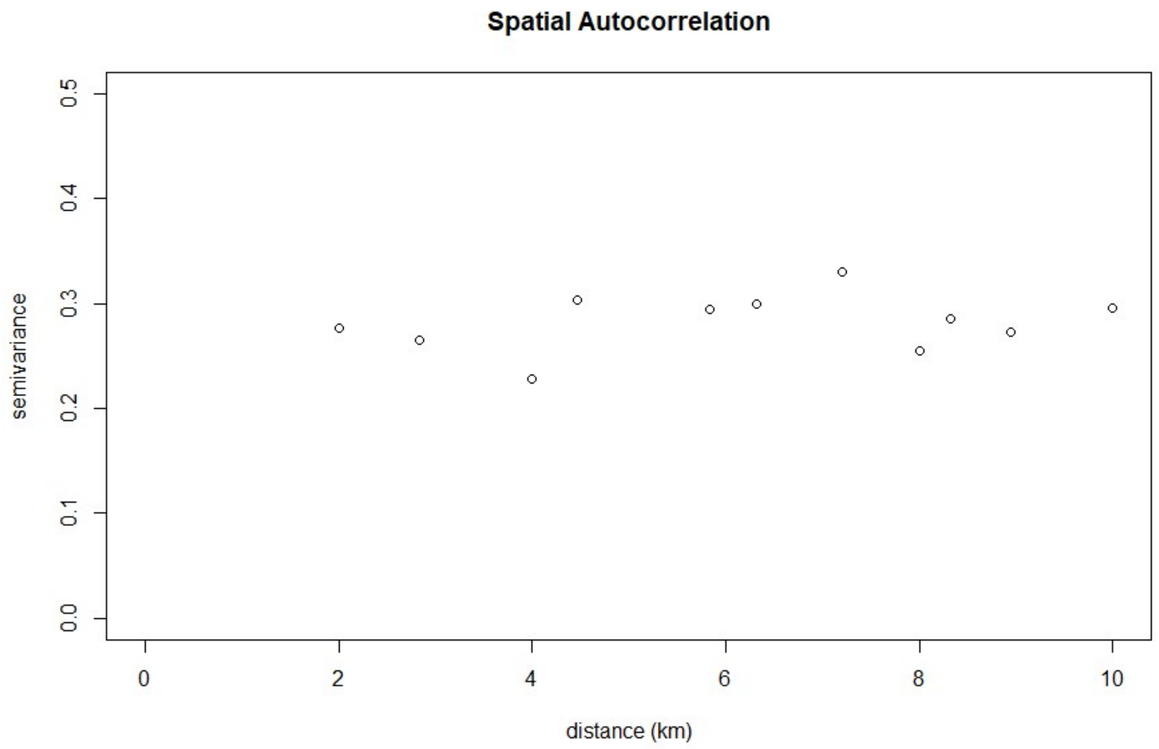


**Fig. S1:** Spatial autocorrelation variogram for the bark stripping model. There is an absence of spatial autocorrelation (no nugget effect). Please note that no sampling stations in the database are closer than 2 km.

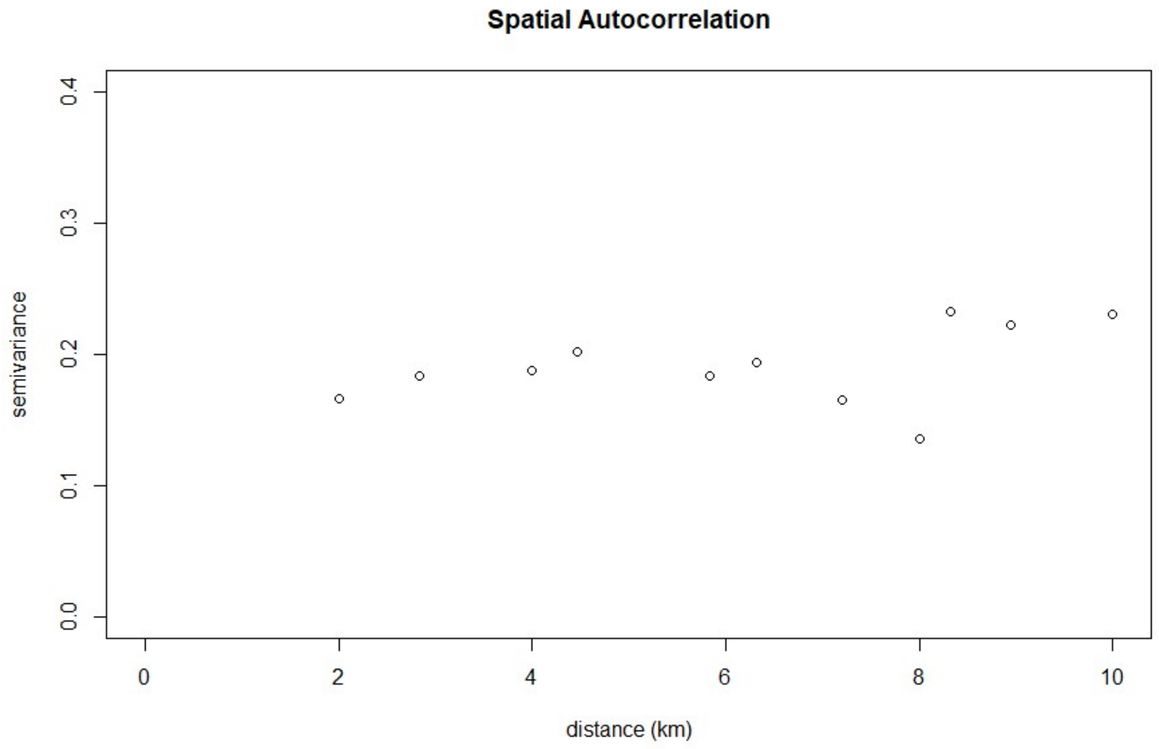


**Fig. S2:** Spatial autocorrelation variogram for the browsing model. There is an absence of spatial autocorrelation (no nugget effect). Please note that no sampling stations in the database are closer than 2 km.

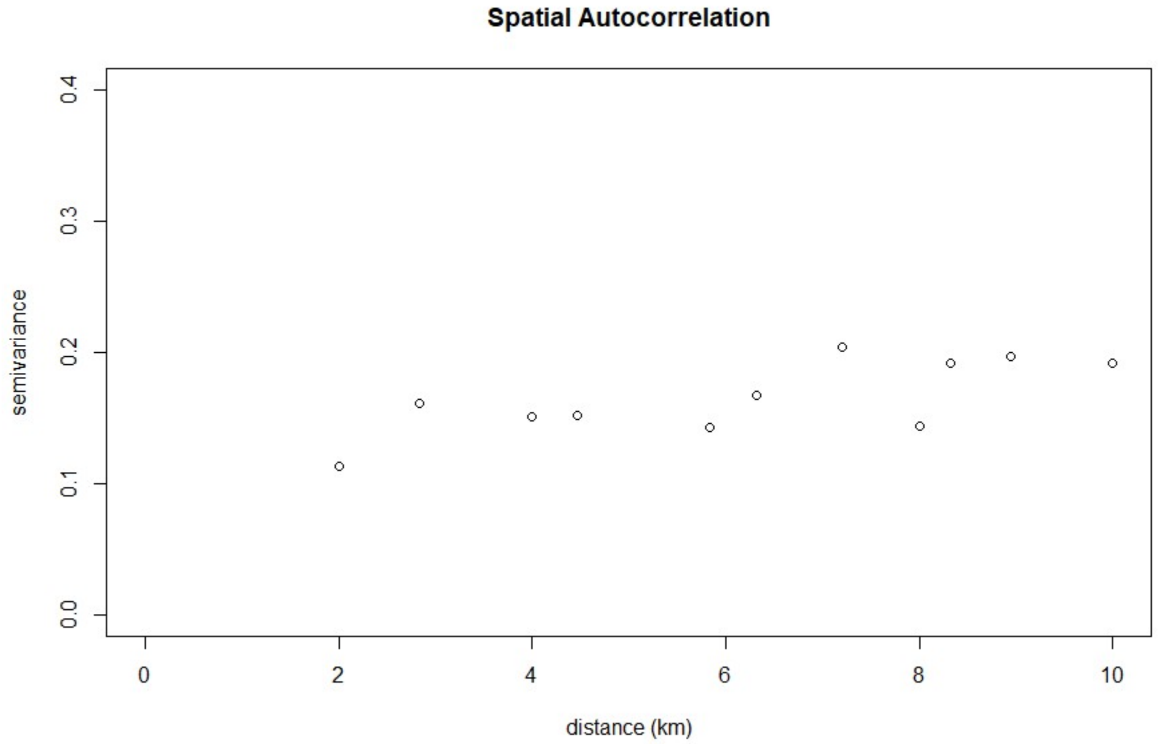


**Fig. S3:** Spatial autocorrelation variogram for fraying model. There is an absence of spatial autocorrelation (no nugget effect). Please note that no sampling stations in the database are closer than 2 km.

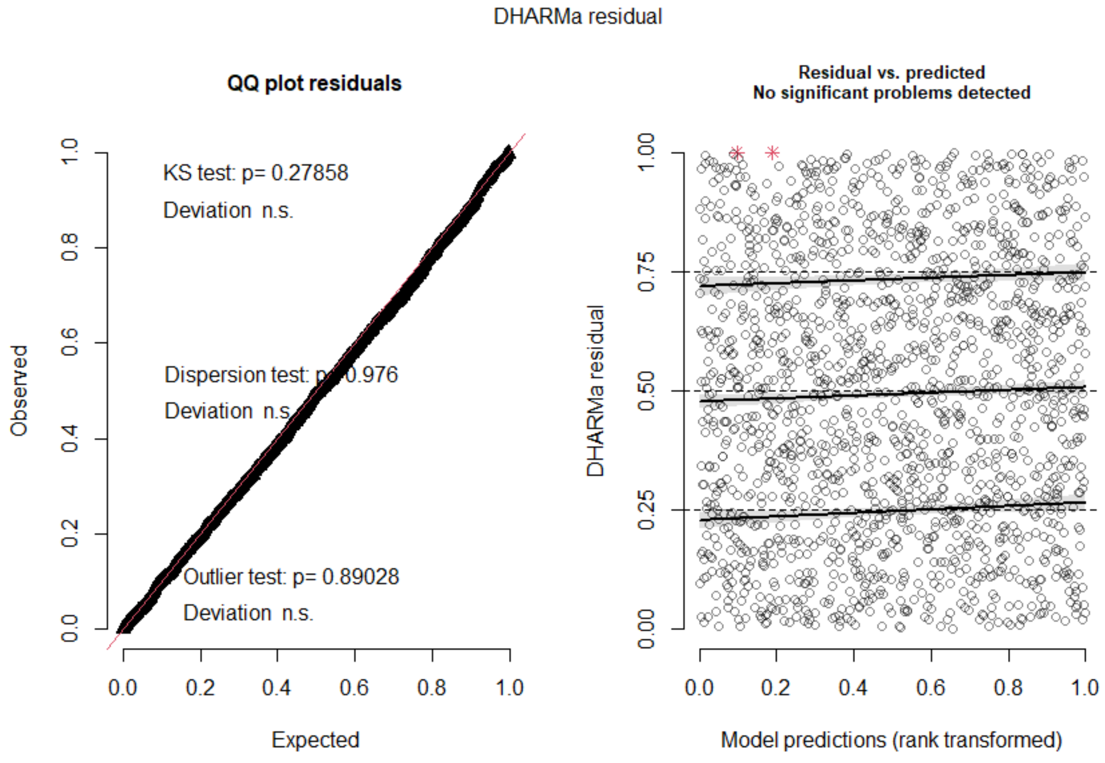


**Fig. S4:** Successful DHARMa model diagnostics for the bark stripping model.

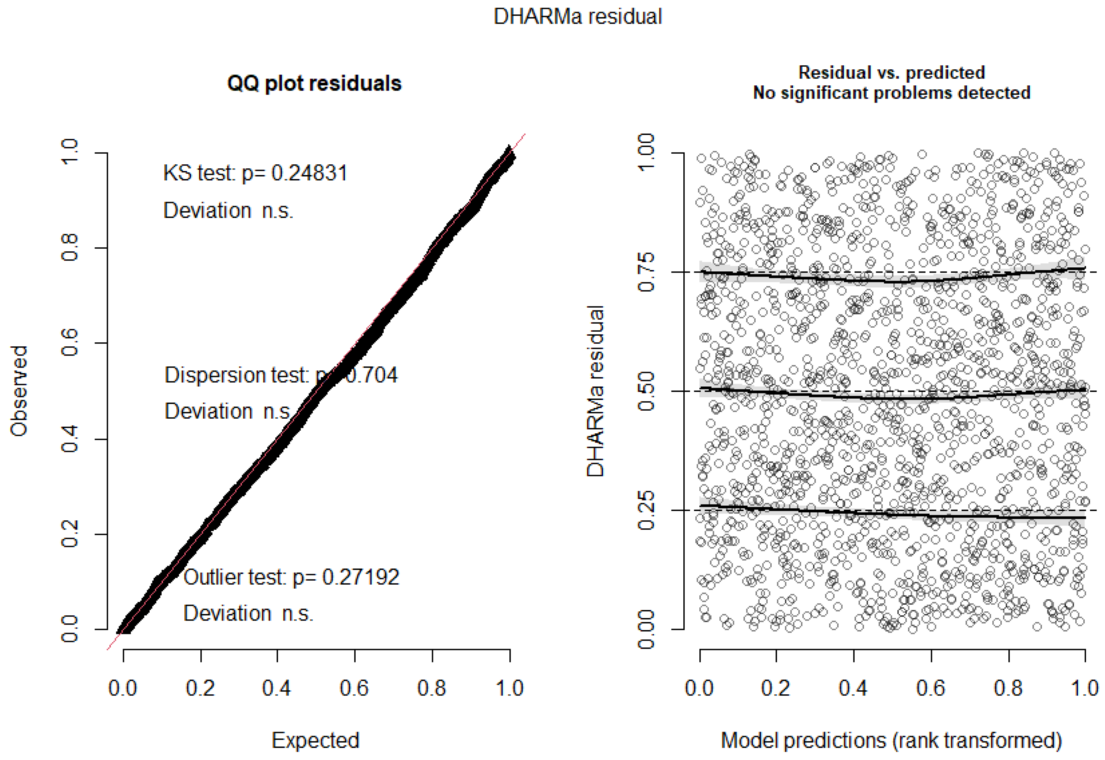


**Fig. S5:** Successful DHARMa model diagnostics for the browsing model.

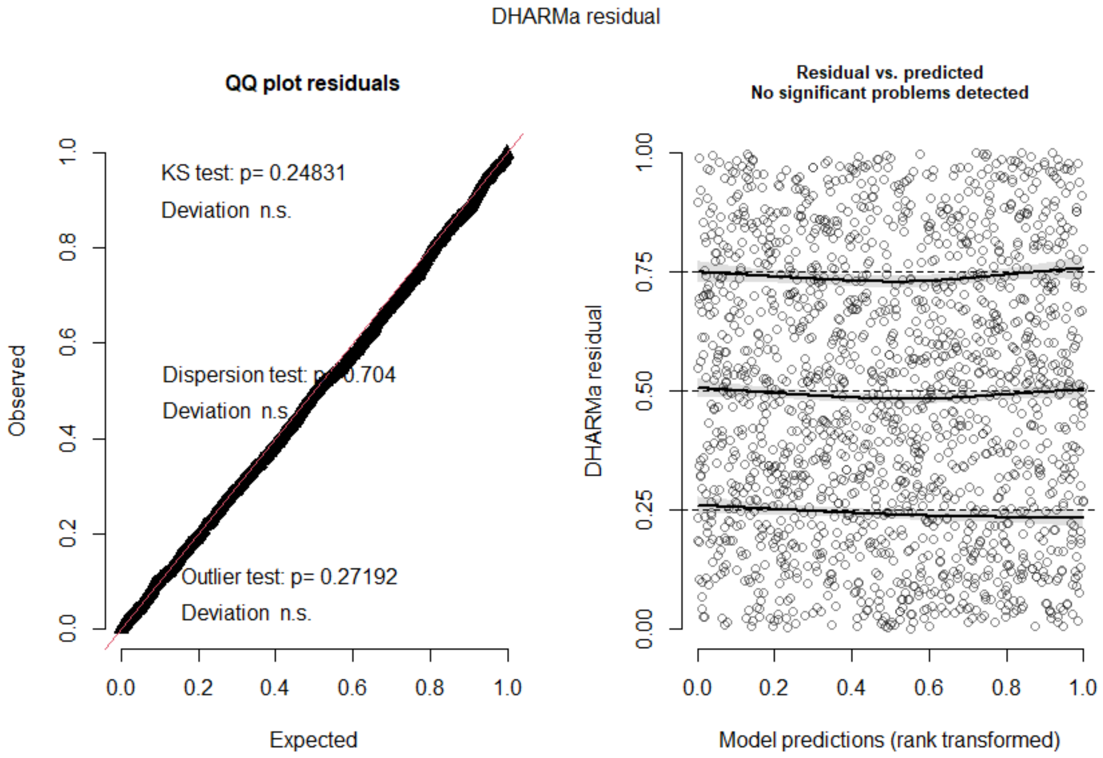


**Fig. S6:** Successful DHARMa model diagnostics for the fraying model.

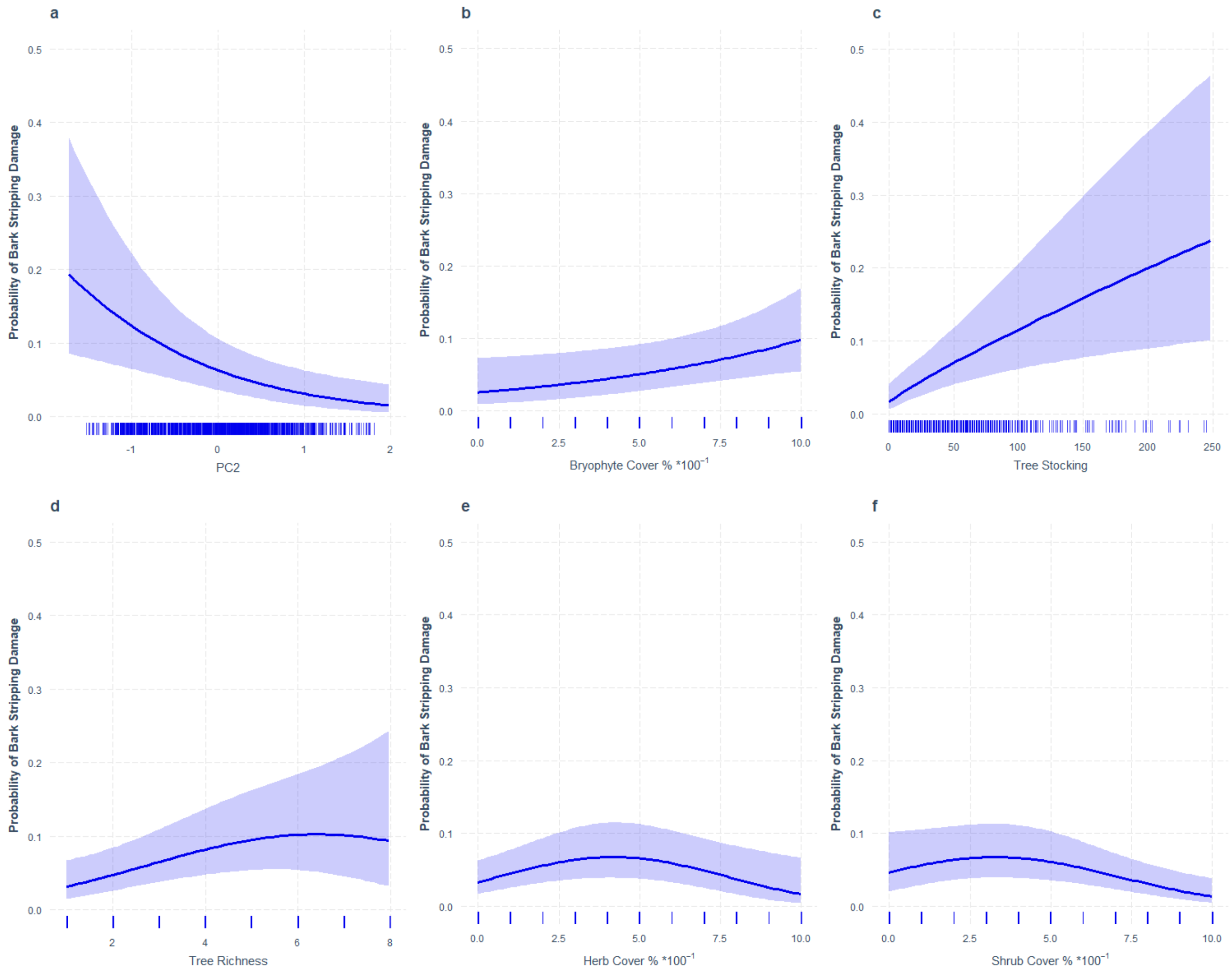


**Fig. S7:** Effect plots for bark stripping damage likelihood predicted by the generalized linear model as a function of a) PC2 values (PC2 increases with decreasing average tree age and increasing standard deviation (SD), thus classifying the forests in a spectrum from mature monoculture (high average age & low SD age) to young natural woodland (low average age & high SD age)), b) bryophyte cover % c) tree stocking, d) tree richness , e) herb cover % and f) shrub cover %. Shaded areas are marginal 95% confidence intervals.

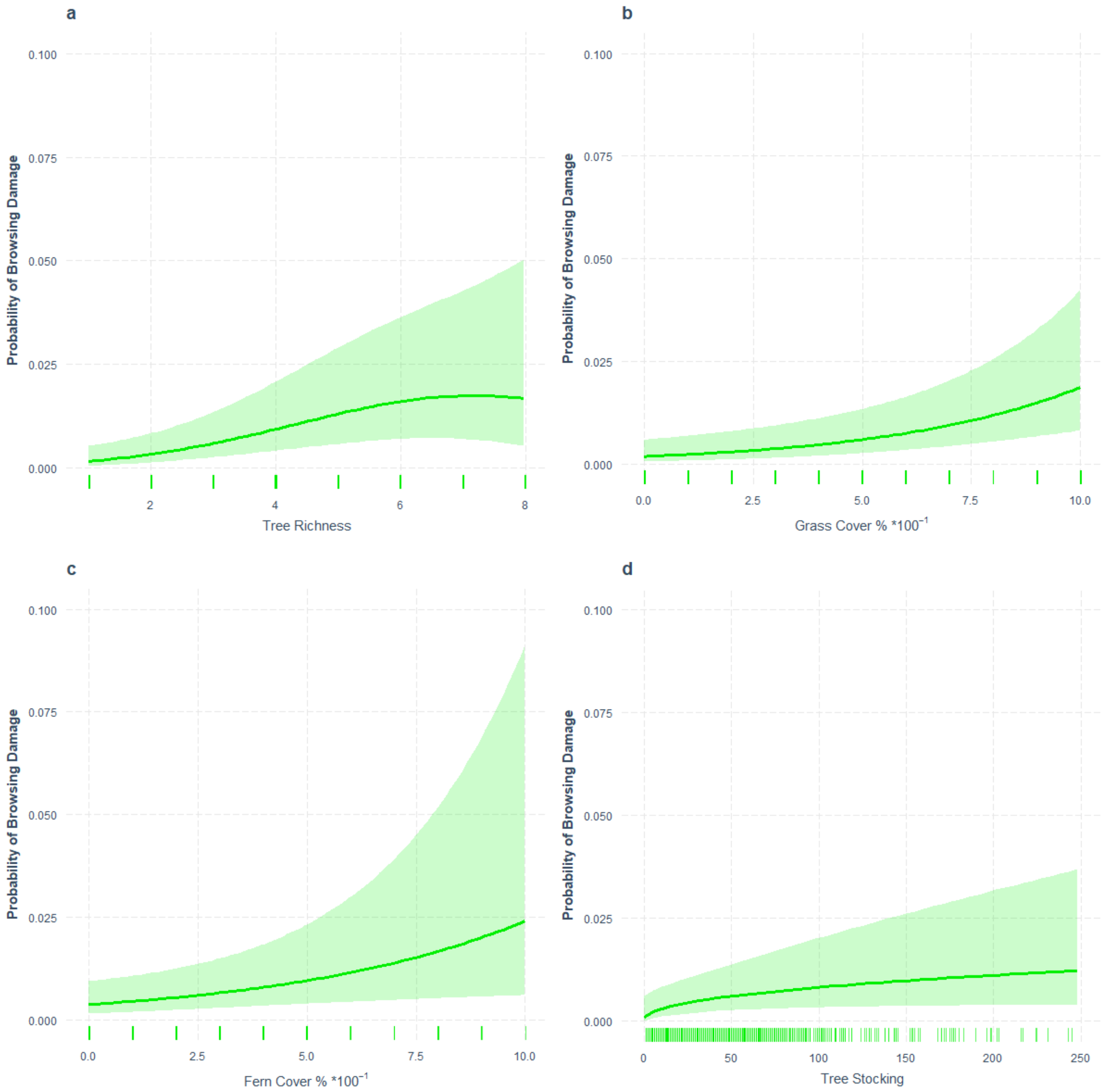


**Fig. S8:** Effect plots for browsing damage likelihood predicted by the generalized linear model as a function of a) tree species richness, b) grass cover %, c) fern cover % and d) tree stocking number. European forest type. Shaded areas are marginal 95% confidence intervals.

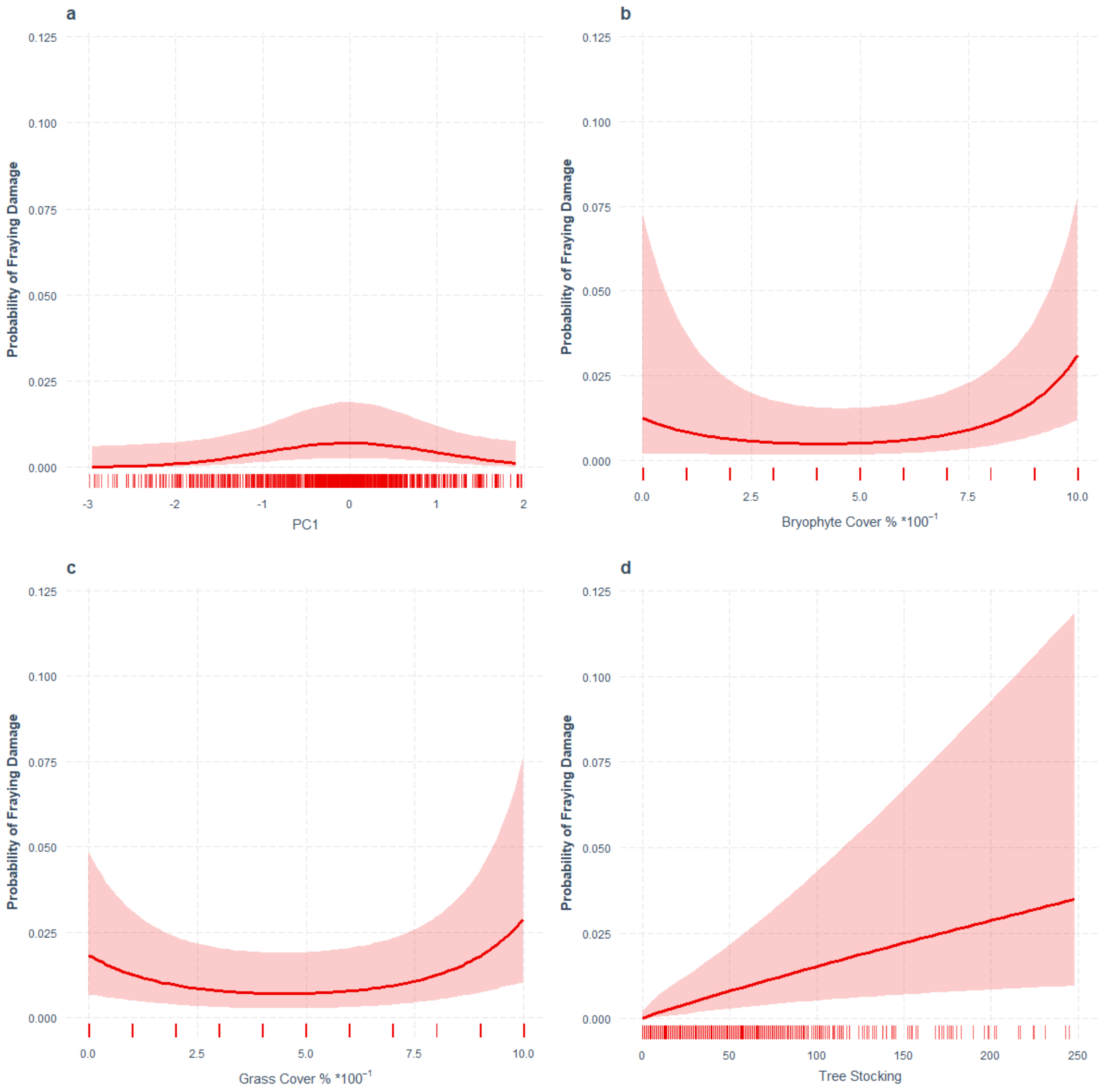


**Fig. S9:** Effect plots for fraying damage likelihood predicted by the generalized linear model as a function of a) PC1 values (PC1 increases with decreasing average tree age and decreasing standard deviation (SD), thus classifying the forests in a spectrum from mature natural woodlands (high average age & high SD age) to young monocultures (low average age & low SD age)), b) bryophyte cover %, c) grass cover % and d) tree stocking number. Shaded areas are marginal 95% confidence intervals.

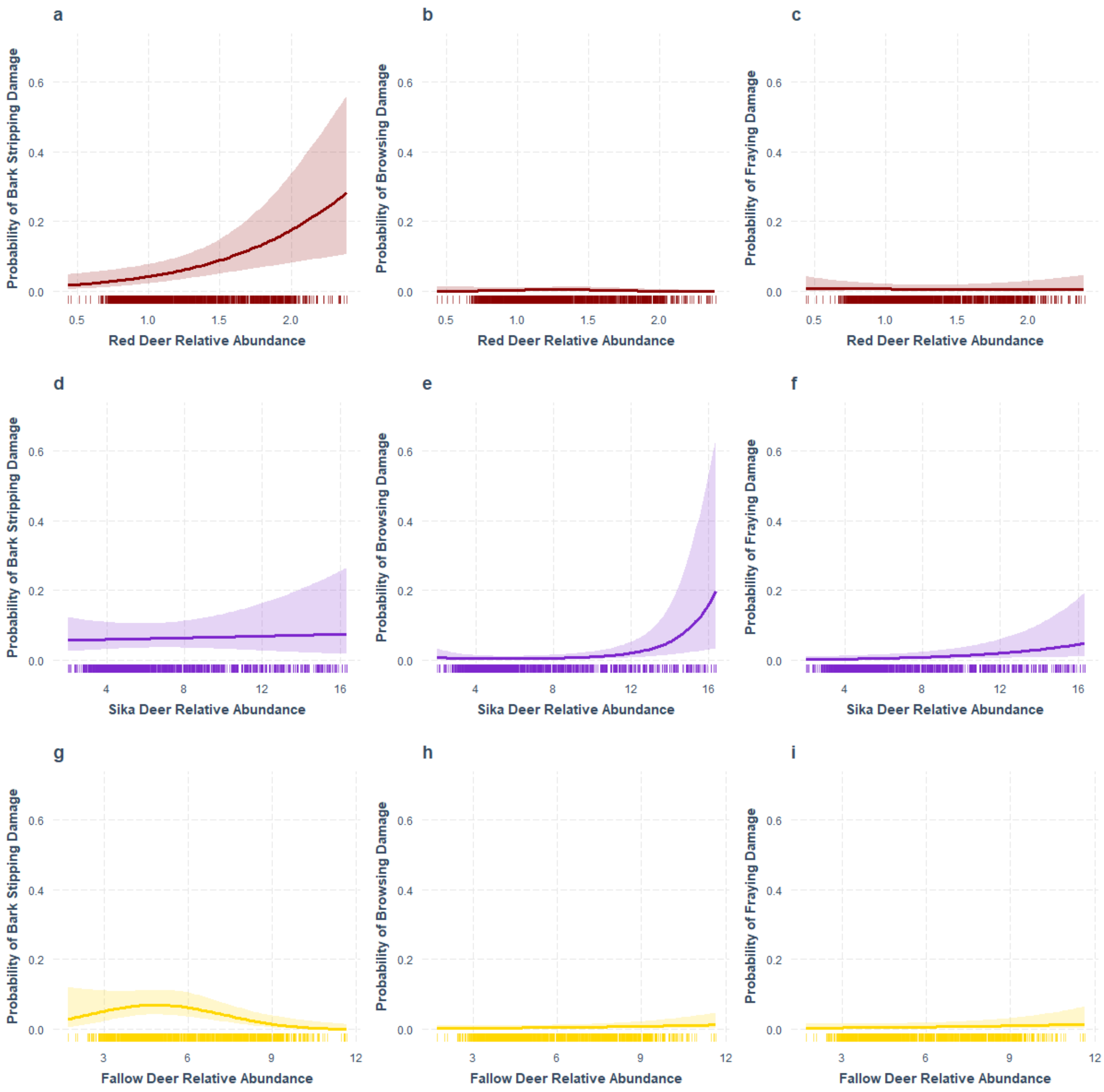


**Fig. S10:** Effect plots for bark stripping damage (left column), browsing damage (middle column) and fraying damage (right column) likelihood predicted by the generalized linear models as a function of a - c) red deer relative abundance, d-f) sika deer relative abundance, & g-i) fallow deer relative abundance. Shaded areas are marginal 95% confidence intervals.

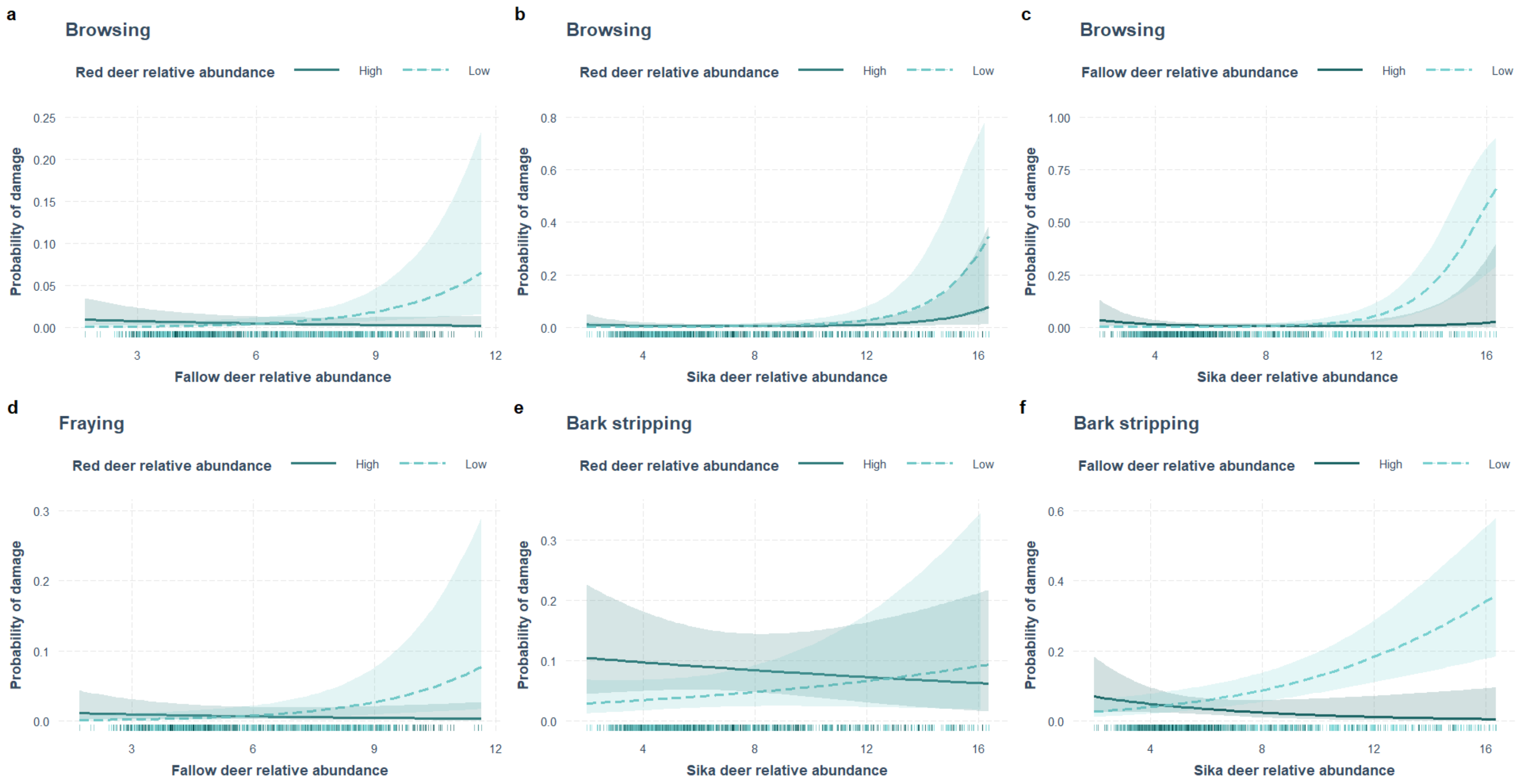


**Fig. S11:** Multiline interaction effect plots predicted by the generalized linear models for: a) browsing damage for red and fallow deer, b) browsing damage for red and sika deer, c) browsing damage for sika and fallow deer, d) fraying damage for red and fallow deer, e) bark stripping damage for red and sika deer and, f) bark stripping damage for sika and fallow deer. Each effect plot shows the respective damage likelihood on the y-axis, increasing deer relative abundance of one species on the x-axis against two fixed values of another species relative abundance, based on the 1$^{st}$ and 3$^{rd}$ Quantile of their respective relative abundance.
